# Dissociable neural mechanisms track evidence accumulation for selection of attention versus action

**DOI:** 10.1101/171454

**Authors:** Amitai Shenhav, Mark A. Straccia, Jonathan D. Cohen, Matthew M. Botvinick

## Abstract

Decision-making is typically studied as a sequential process from the selection of what to attend (e.g., between possible tasks, stimuli, or stimulus attributes) to the selection of which actions to take based on the attended information. However, people often gather information across these levels in parallel. For instance, even as they choose their actions, they may continue to evaluate how much to attend other tasks or dimensions of information within a task. We scanned participants while they made such parallel evaluations, simultaneously weighing how much to attend two dynamic stimulus attributes and which response to give based on the attended information. Regions of prefrontal cortex tracked information about the stimulus attributes in dissociable ways, related to either the predicted reward (ventromedial prefrontal cortex) or the degree to which that attribute was being attended (dorsal anterior cingulate, dACC). Within dACC, adjacent regions tracked uncertainty at different levels of the decision, regarding what to attend versus how to respond. These findings bridge research on perceptual and value-based decision-making, demonstrating that people dynamically integrate information in parallel across different levels of decision making.

Naturalistic decisions allow an individual to weigh their options within a particular task (e.g., how best to word the introduction to a paper) while also weighing how much to attend other tasks (e.g., responding to e-mails). These different types of decision-making have a hierarchical but reciprocal relationship: Decisions at higher levels inform the focus of attention at lower levels (e.g., whether to select between citations or email addresses) while, at the same time, information at lower levels (e.g., the salience of an incoming email) informs decisions regarding which task to attend. Critically, recent studies suggest that decisions across these levels may occur in parallel, continuously informed by information that is integrated from the environment and from one’s internal milieu^1,2^.

Research on cognitive control and perceptual decision-making has examined how responses are selected when attentional targets are clearly defined (e.g., based on instruction to attend a stimulus dimension), including cases in which responding requires accumulating information regarding a noisy percept (e.g., evidence favoring a left or right response)^3-7^. Separate research on value-based decision-making has examined how individuals select which stimulus dimension(s) to attend in order to maximize their expected rewards^8-11^. However, it remains unclear how the accumulation of evidence to select high-level goals and/or attentional targets interacts with the simultaneous accumulation of evidence to select responses according to those goals (e.g., based on the perceptual properties of the stimuli). Recent work has highlighted the importance of such interactions to understanding task selection^12-15^, multi-attribute decision-making^16-18^, foraging behavior^19-21^, cognitive effort^22,23^, and self-control^24-27^.

While these interactions remain poorly understood, previous research has identified candidate neural mechanisms associated with multi-attribute value-based decision-making^11,28,29^ and with selecting a response based on noisy information from an instructed attentional target^3–5^. These research areas have implicated the ventromedial prefrontal cortex (vmPFC) in tracking the value of potential targets of attention (e.g., stimulus attributes)^8,11^ and the dorsal anterior cingulate cortex (dACC) in tracking an individual’s uncertainty regarding which response to select^30–32^. It has been further proposed that dACC may differentiate between uncertainty at each of these parallel levels of decision-making (e.g., at the level of task goals or strategies vs. specific motor actions), and that these may be separately encoded at different locations along the dACC’s rostrocaudal axis^32,33^. However, neural activity within and across these prefrontal regions has not yet been examined in a setting in which information is weighed at both levels within and across trials.

Here we use a value-based perceptual decision-making task to examine how people integrate different dynamic sources of information to decide (a) which perceptual attribute to attend and (b) how to respond based on the evidence for that attribute. Participants performed a task in which they regularly faced a conflict between attending the stimulus attribute that offered the greater reward or the attribute that was more perceptually salient (akin to persevering in writing one’s paper when an enticing email awaits). We demonstrate that dACC and vmPFC track evidence for the two attributes in dissociable ways. Across these regions, vmPFC weighs attribute evidence by the reward it predicts and dACC weighs it by its attentional priority (i.e., the degree to which that attribute drives choice). Within dACC, adjacent regions differentiated between uncertainty at the two levels of the decision, regarding what to attend (rostral dACC) versus how to respond (caudal dACC).

## Results

Participants were shown random dot kinematograms that varied along two dimensions, direction of dot motion (up or down) and the dominant dot color (blue or red) (Fig. 1)^3,4^. They gave a single response on each trial (left or right), which could be correct for neither, one, or both attributes (Fig. 1A). Participants were allowed to freely choose how much to rely on each attribute in selecting their response, and were rewarded for each attribute they responded to correctly. We independently varied the level of perceptual noise (i.e., the discriminability) of the two attributes across trials, such that motion or color information could be more salient on a given trial (Fig. 1B). Correct responses for the two attributes were either rewarded equally (Epoch 1) or one attribute was rewarded twice as much as the other (Epochs 2-3) (Fig. 1C), biasing attention towards the more rewarded attribute on a given block of trials.

**Figure 1.**
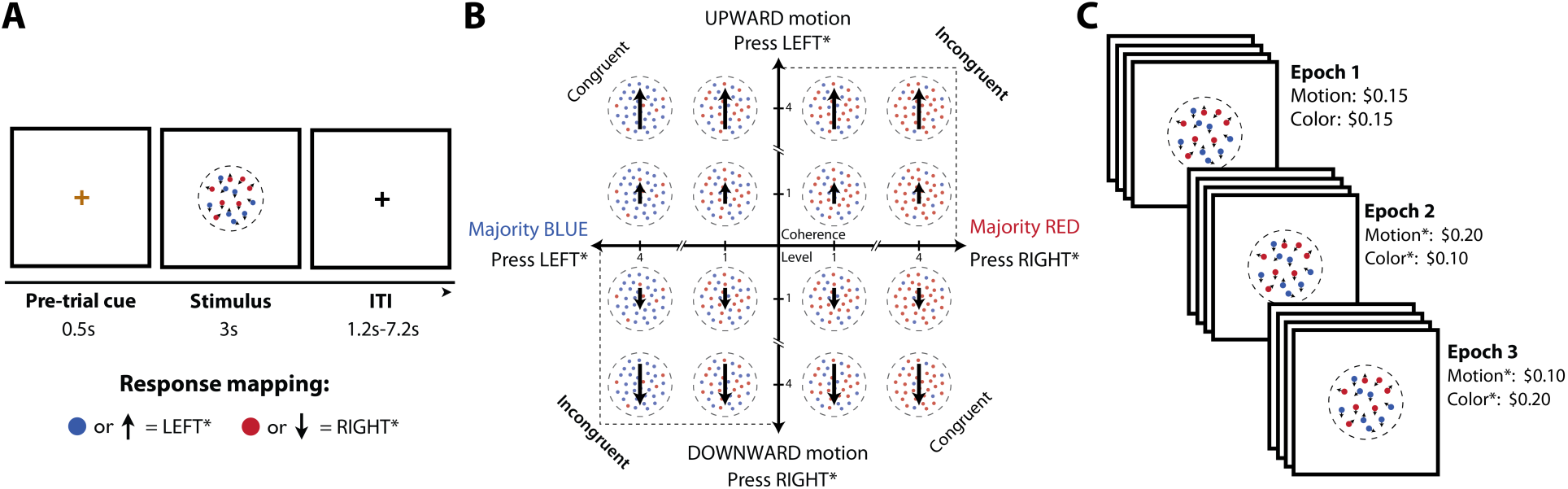
Behavioral paradigm. **A)** Participants viewed random dot motion patterns and could indicate whether the dots were primarily moving up or down and/or whether they were majority red or blue. They responded with either a left or right button press. Responses were bivalent, denoting both a color and a motion direction, and participants were rewarded based on the number of dimensions for which this response was correct. **B)** The coherence and correct response for motion and color dimensions were varied orthogonally across trials. Four participant-specific coherence levels were used for each attribute. ^∗^Response mappings and Epoch 2-3 reward associations were counter-balanced across participants. **C)** Participants performed three epochs (192 trials each) that varied in motion/color reward associations, either rewarding both equally (Epoch 1) or differently (Epochs 2-3).

### Effect of attribute evidence on choice

To examine the influence of each stimulus attribute on choice, we entered the coherence of the two attributes into a mixed-effects logistic regression predicting choice on a given trial. Focusing first on the initial task epoch – during which the two attributes were rewarded equally – we found that, as expected, subjects’ choices were significantly influenced by the coherence of both motion (b = 1.5, SE = 0.11, *z* = 13.7) and color (b = 0.9, SE = 0.09, *z* = 10.1, *p*s < 0.0001). The more evidence provided by either attribute in favor of a given response, the more likely that response. Overall, choices were also more influenced by motion than color evidence in this initial (baseline) block (b = 0.39, SE = 0.11, *t* = 3.7, *p* < 0.001).

In Epochs 2-3 of the session, correct responses for one attribute were more highly rewarded than the other (either motion or color, counter-balanced across segments and participants). During these epochs we found that subjects weighed their decisions much more heavily toward the more rewarding attribute (b = 1.9, SE= 0.13, *t* =14.3) but the low reward attribute continued to exert a significant influence (b = 0.45, SE= 0.06, *t* = 7.8, *p*s < 0.0001; Fig. 2). RTs were also faster the more evidence supported the chosen response (high-reward: b = −0.34, SE = 0.03 *t* = −11.9; low-reward: b = −0.05, SE = 0.01 *t* = −5.6, *p*s < 0.001). Effects of both high and low reward attributes on choices and RTs held irrespective of whether motion or color was more highly rewarded (*t*s > 3.5, *p*s < 0.002).

**Figure 2.**
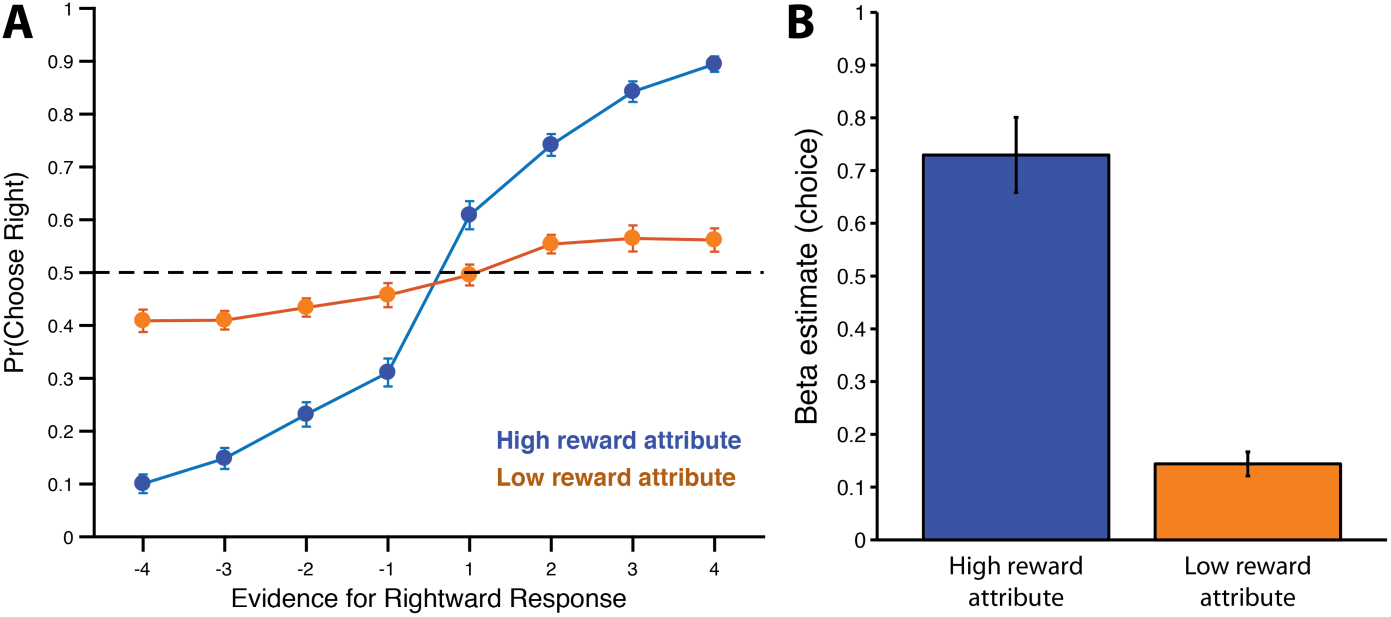
Behavioral sensitivity to attribute evidence and rewards. During Epochs 2-3, responses were highly sensitive to both evidence (coherence) and the relative reward for the two attributes. **A)** A psychometric curve shows that participants were much more likely to select a response the more evidence it provided for the high-reward attribute. **B)** Average regression coefficients for the influence of high and low reward coherence on choice. While high-reward attribute coherence exerted the strongest influence on responses, participants were still sensitive to the evidence supporting the low-reward attribute. Error bars reflect s.e.m.

### Dissociable correlates of attribute evidence in dACC and vmPFC

Given their previous involvement in evidence integration for perceptual and/or value-based decisions, we tested the degree to which dACC and vmPFC tracked the perceptual evidence supporting the chosen response (e.g., if the left response was made on a given trial, this is the signed motion and color coherence level in support of the left response).

Consistent with previous findings^3,11,16,34,35^, we found that vmPFC tracked how much total evidence was available for the chosen option (b = 0.05, SE = 0.01, *t_vmPFC_* = 3.7,*p* < 0.001) while dACC tracked how little evidence was available for that option (b = −0.07, SE = 0.01, *t_dACC_* = −6.8, *p* < 0.001). Furthermore, consistent with previous studies of value-based integration of stimulus attributes^11,16,36,37^, we found that vmPFC encoded the evidence favoring the chosen option from both the higher reward attribute (b = 0.055, SE = 0.015, *t* = 3.7,*p* < 0.001) and the lower reward attribute (b = 0.03, SE = 0.01, *t* = 2.3,*p* < 0.03). The vmPFC’s relative encoding of evidence for these two attributes was in fact almost identical to the relative reward provided for a correct response along these attributes (vmPFC ratio: 1.99:1; actual ratio: 2:1) (Fig. 3B). This is particularly notable given that participants were not given trial-wise feedback about their performance. By contrast, dACC was primarily sensitive to the (inverse) evidence for the high-reward attribute (b = −0.11, SE = 0.02, *t* = −7.3, *p* < 0.001) and exhibited a weaker and non-significant trend for evidence of the low-reward attribute (b = −0.01, SE = 0.01, *t* = −1.6, *p* = 0.13) (Fig. 3C).

**Figure 3.**
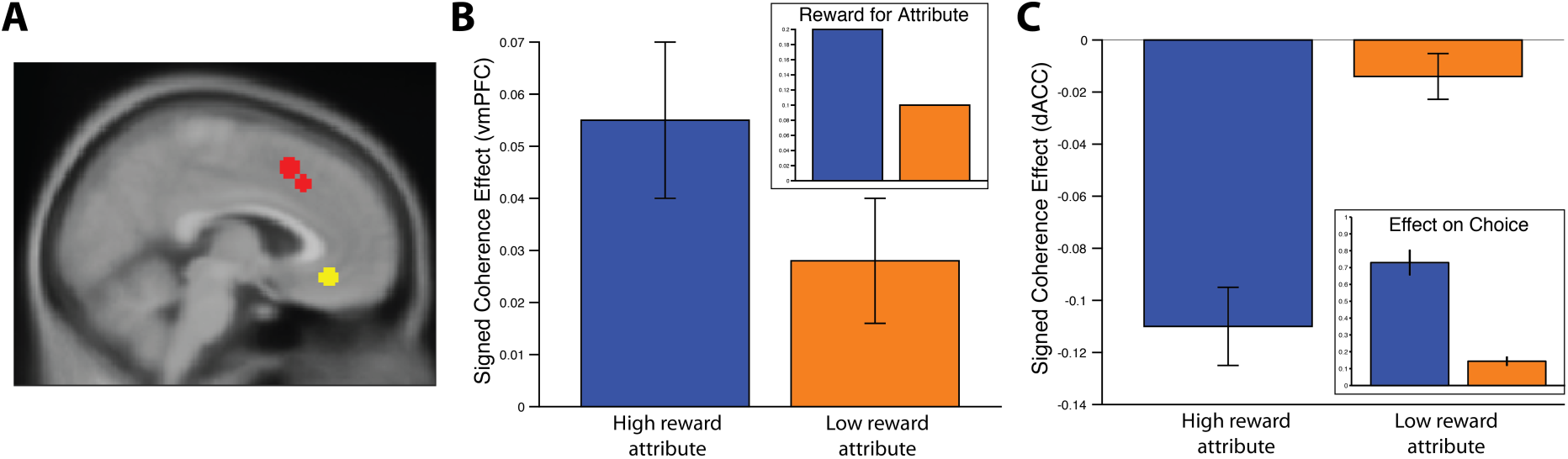
vmPFC and dACC differentially encode evidence for the high and low reward attributes. **A)** vmPFC (yellow) and dACC (red) ROIs were defined a priori based on relevant findings from research on integration of information from multi-attribute stimuli^3,11^. **B)** vmPFC positively tracked the evidence each attribute provided for the chosen response (signed coherence), but it did not weigh evidence for both attributes equally. Rather, responses to the two attributes were weighed in proportion to the reward expected for responding correctly to that attribute. For reference, the inset shows the reward amounts (in dollars) expected for each attribute. **C)** dACC tracked how little evidence was available for these two attributes, weighing evidence for the two attributes in proportion to the influence that attribute will have on the ultimate choice (inset from Fig. 2B), potentially reflecting the amount of attention placed on that attribute while forming a decision. Error bars reflect s.e.m.

As explored further below, these findings tentatively suggest that vmPFC signals of attribute evidence scale with the expected reward for that attribute (compare Fig. 3B inset) whereas equivalent signals of attribute evidence in dACC scale with the influence that attribute has on the ultimate decision (and therefore how much attention was likely paid to that attribute prior to making a decision) (compare Fig. 3C inset). Accordingly, vmPFC activity was greater on trials where motion and color information supported the same response (b = 0.05, SE = 0.02, *t* = 2.5, *p* < 0.02) while dACC, with its primary emphasis on the high-reward attribute, did not encode whether the alternate attribute provided congruent information (*t* = −0.39, *p* > 0.70). This region of dACC was therefore sensitive to uncertainty at the level of which response to give (left or right) but only as it pertained to the more rewarding attribute. Follow-up analyses found comparable effects in regions that were independently identified as being most sensitive to color and motion (Supplementary Analysis 1).

These findings demonstrate the degree to which these two regions respond to evidence for the response that was chosen on a given trial. As such, they point to potential roles these regions may play during decision-making about which action to select. In order to instead examine the role these regions may play in higher-level decisions about which attribute to attend, we can instead examine the degree to which these regions track the salience of each attribute (i.e., how much evidence was available for a given attribute, irrespective of the response it supported; also referred to as its *unsigned* coherence). In particular, given that participants heavily prioritized the high-reward attribute when selecting their ultimate response (Fig. 2), increased salience of that attribute might serve to increase their confidence in the decision to focus their attention on it. Conversely, as the salience of the low-reward attribute increases, the participant may experience greater uncertainty about whether to continue focusing on the high-reward attribute or whether to instead focus more on the low-reward attribute.

We found a significant interaction in the degree to which these two regions tracked the salience of the high versus low reward attributes: dACC negatively tracked the salience of the high reward attribute and positively tracked the salience of the low reward attribute, while vmPFC showed the reverse pattern (Supplementary Fig. 1; dACC vs. vmPFC: b*_high_* = −0.07, SE*_high_* = 0.01, *t_high_* = −6.5, *p* < 0.001, b_*low*_ = 0.02, SE*_low_* = 0.01, *t_low_* = 2.4, *p* < 0.02).

### Encoding of levels of uncertainty along dACC’s rostrocaudal axis

The previous analyses demonstrated that dACC and vmPFC tracked the salience of the low-reward attribute with opposite signs. While they provide preliminary evidence that the salience of the low-reward attribute may be tracked negatively in vmPFC and positively in dACC, these effects of low-reward attribute salience were individually non-significant. However, given that our a priori dACC ROI was based on the dACC’s response in a previous study^3^ to the attribute an individual was instructed to attend – which may correspond to the high-reward attribute in the current study – we considered the possibility that this may not have been the optimal choice of ROI for capturing a reliable effect of the low-reward attribute. Therefore, we performed a whole-brain analysis to examine whether responses to attribute salience varied outside of this region of dACC. When doing so, we found a striking distinction: whereas a more caudal region of dACC was sensitive to the *absence* of evidence for the high reward attribute, a more rostral region was sensitive to the *availability* of evidence for the low reward attribute (Fig. 4A).

**Figure 4.**
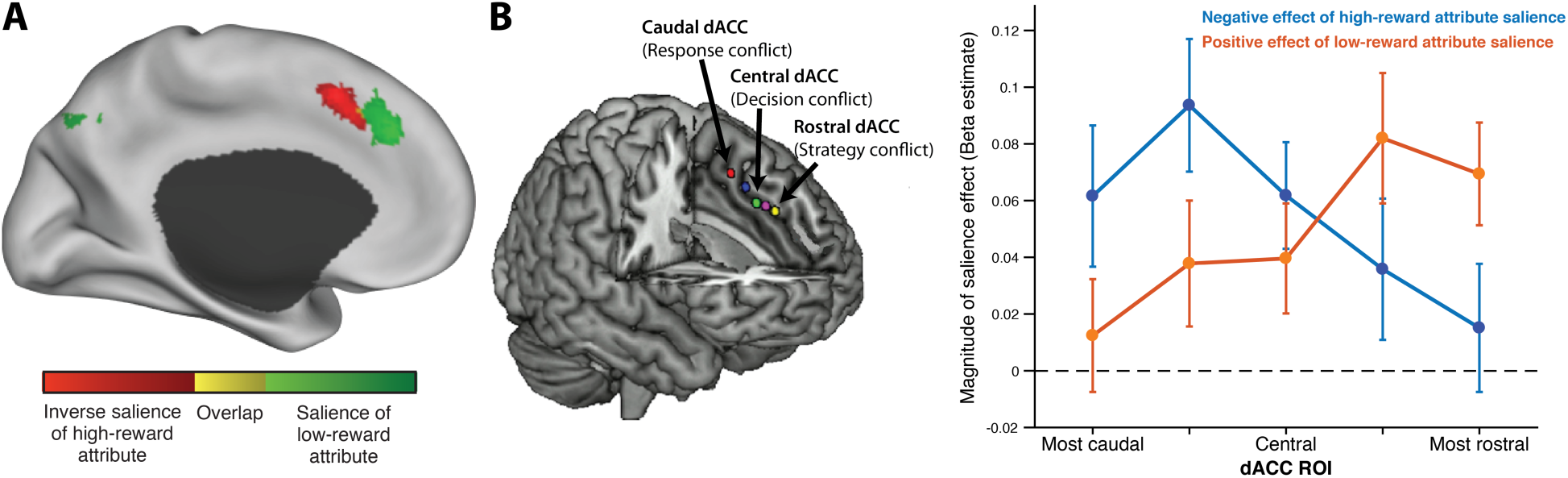
dACC encoding of attribute salience varies along rostrocaudal axis. **A)** A whole-brain analysis revealed that more caudal regions of dACC negatively tracked the salience of the high-reward attribute (red) and more rostral regions positively tracked the salience of the low-reward attribute (green). Activations reflect *t*-statistics, whole-brain corrected to achieve a clusterwise *p*<0.05. **B)** This rostocaudal pattern was confirmed with average beta estimates extracted from a set of independent ROIs drawn from an earlier study^33^, which proposed that these reflect a range of uncertainty/conflict levels, from low-level responses (e.g., motor actions) most caudally to more abstract responses (e.g., decisions and strategies) more rostrally. Left panel modified from Ref. ^33^. Error bars reflect s.e.m.

While not initially expected, this anatomical distinction appeared to be consistent with previous proposals that signals of cognitive demand may be topographically organized along a rostrocaudal axis within dACC^33,38^. This work has shown that increasingly rostral regions of dACC track cognitive demands related to increasingly abstract or complex control targets – ranging from uncertainty/conflict between potential motor actions (caudal-most) to potential decision options (central) to potential strategies (rostral-most) – paralleling similar patterns of representational abstraction on the lateral surface^39,40^. We therefore sought to test whether the anatomical distinction we observed reflected a functional dissociation along this proposed rostrocaudal axis, between uncertainty at the level of responses (left vs. right) and uncertainty at the level of attentional targets (motion vs. color attribute).

Analyses along this rostrocaudal axis confirmed the presence of such a dissociation (Fig. 4B): high reward salience is negatively tracked in more caudal ROIs and low reward salience is positively tracked in more rostral ROIs. To test this dissociation more explicitly, we compared salience encoding in the two most rostral and the two most caudal ROIs. This analysis revealed a significant interaction between ROI location and type of salience encoding, with the rostral ROIs tracking the salience of the low reward attribute more positively than the caudal ROIs (b = 0.03, SE = 0.01, *t* = 3.1, *p* < 0.005) and the caudal ROIs tracking the salience of the high-reward attribute more negatively than the rostral ROIs (b = 0.03, SE = 0.01, *t* = 2.4, *p* < 0.03). Moreover, the effect of low reward attribute salience in the rostral-most ROIs remained even when restricting our analysis to trials on which motion and color evidence supported the same response (congruent trials; b = 0.05, SE = 0.01, *t* = 3.5, *p*<0.001), suggesting that these regions were not simply tracking whether the low-reward attribute was strongly supporting a different response than the high reward attribute^41^.

Since regions of dACC have been previously implicated in encoding other signals of cognitive demand, including various forms of errors^41–44^, we further tested the extent to which each of these regions also signaled errors in the current task. In addition to tracking the salience of these attributes, these regions of dACC also encoded whether the participant committed a high-reward error on a given trial (caudal: b = 0.16, SE = 0.03, *t* = 5.6; rostral: b = 0.17, SE = 0.03, *t* = 4.9, *p*s<0.001). While we excluded missed trials from all other analyses, when specifically including this as a regressor we also find, as expected, that both regions of dACC also exhibit increased activity when participants fail to respond by the deadline (caudal: b = 0.81, SE = 0.08, *t* = 9.7; rostral: b = 0.45, SE = 0.07, *t* = 6.3, *p*<0.001), independent of the salience of the attributes. Caudal ROIs responded more strongly than rostral ROIs to having missed a response (b = 0.38, SE = 0.09, *t* = 4.4, *p* < 0.001).

### Rostral dACC and attention to low-reward attribute salience

We found that rostral dACC positively encoded the salience of the low-reward attribute, which competes with the high-reward attribute for attention and influence over the ultimate decision. We therefore performed logistic regressions to test whether activity in this region affected how much influence the low-reward attribute exerted on choice. We found that the likelihood of providing the correct response for the low-reward attribute was predicted by an interaction between rostral dACC activity and the salience of that attribute (b = 0.05, SE = 0.02, *z* = 2.6, *p* < 0.02). In other words, increased activity in this region was associated with an increased likelihood that the participant responded according to a salient low-reward attribute. This was not true for the interaction of rostral dACC with high-reward attribute salience (*z* = 0.36, *p* = 0.72), nor was it true for the interaction of caudal dACC with low-reward attribute salience (*z* = 1.3, *p* = 0.18).

### Trial history effects in vmPFC

We performed a final exploratory analysis to examine whether activity in vmPFC and/or dACC reflected evidence accumulated not only on the current trial (as reported above) but also from previous trials. We found this to be the case in vmPFC – controlling for the signed coherence of the two attributes on the current trial, activity in vmPFC was greater when more high-reward attribute information had been available to support the response chosen on the last trial (b = 0.03, SE = 0.01, *t* = 2.5, *p*<0.02). There was no significant effect of the previous signed coherence of the low-reward attribute (*t* = −0.31, *p*=0.76), nor did either region of dACC track the previous signed coherence of either attribute (|*t*s| < 1.0, *p*s>0.30).

## Discussion

Most everyday tasks invoke a natural tension between focusing on the current task and switching to an alternative. Rather than committing to a given task and then performing it, individuals typically face a recurring decision regarding which task to attend and how much^22,23,31^. In the current study, we asked participants to make perceptual decisions involving two parallel streams of visual evidence (motion direction and color proportion), allowing them to select how much to allow each stream to guide their choice. As a result, their decisions were two-fold: (1) how much to attend each stream and (2) which motor response to select. Whereas the latter decision was influenced by the overall evidence in favor of each response (i.e., upward vs. downward motion, concentration of blue vs. red), the former was influenced by the available reward and the salience (absolute coherence) of a given attribute. We found that the uncertainty associated with each of these two decisions was encoded in adjacent but distinct regions of dorsal ACC: a more caudal region tracked the uncertainty in discriminating evidence for a left versus right response (replicating previous findings^3,30,45^), while a more anterior region appeared to track the uncertainty in selecting which attribute to attend. Specifically, anterior dACC increased activity with the relative ease of attending the less preferred attribute on a given block.

Our findings within dACC are consistent with previous proposals that this region signals demands for cognitive control (e.g., conflict, error likelihood^46–48^) and that these demands may be differentially encoded across different populations within dACC^41,42^. Most notably, our findings are broadly consistent with the recent proposal that dACC signals such demands in a hierarchical manner^32,33,38^ (cf. Refs.^14,49^). Specifically, it has been suggested that dACC contains a topographic representation of potential control demands, with more caudal regions reflecting demands at the level of individual motor responses and more rostral regions reflecting demands at increasing levels of abstraction (e.g., at the level of effector-agnostic response options). According to this framework, it is reasonable to assume that this rostrocaudal axis might encode uncertainty regarding which attribute to attend more rostrally than uncertainty regarding which response to select. Our findings may also be consistent with a more recent proposal that a similar axis within dACC tracks the likelihood of responses and outcomes (e.g., error likelihood) at similarly increasing levels of abstraction^50^. Collectively these accounts of the current findings are consistent with our theory that regions of dACC integrate information regarding the costs and benefits of control allocation (including traditional signals of control demand) in order to adaptively adjust control allocation^31,48^.

The dACC signals we observed are also consistent with evaluation processes unrelated to control per se, indicating for instance the costs of maintaining the current course of action in caudal dACC and the value of pursuing an alternate course of action (cf. foraging) in rostral dACC^19,20^. The connection between rostral dACC activity and choices to follow evidence for the low-reward attribute can be seen as further support for such an account (though this could similarly reflect adjustments of attentional allocation). Our current study is limited in adjudicating between these two accounts because increasing evidence in support of an alternative attentional target in our task (i.e., increased salience of the low-reward attribute) necessarily leads to greater uncertainty regarding whether to continue to focus on the high-reward attribute. However, given that evidence for foraging-specific value signals in dACC remains inconsistent^35,48,51^, an interpretation of our findings that appeals to cognitive costs or demands may be more parsimonious. That said, future studies are required to substantiate the current interpretation by demonstrating that the dACC’s response to the tempting alternative (the salient low-reward attribute) decreases when the relative salience and reward of the alternate attribute are such that the decision to switch one’s target of attention is easy.

In contrast to dACC, where activity tracked how little evidence was available to support the chosen response (i.e., to discriminate between the correct and incorrect response), vmPFC instead tracked the evidence in favor of the chosen response, in a manner proportional to the reward expected for information about each attribute. This finding is broadly consistent with previous findings in the value-based decision making literature, where vmPFC is often associated with the value of the chosen option and/or its relationship to the value of the unchosen option^52,53^. The fact that vmPFC’s weights on these attributes were *not* proportional to the weight each attribute was given in the final decision suggests that vmPFC may have played less of a role in determining how this information was used to guide a response, than in providing an overall estimate of expected reward. In addition to any incidental influence it may have on the perceptual decision on a given trial, this reward estimate could provide a learning signal about the task context more generally (e.g., overall reward rate)^26^, consistent with our observation that this region encodes elements of reward expected from a previous trial.

Previous research has identified a number of parallels between behavioral and neural patterns evoked by perceptual and value-based decisions^54–56^. Both have been well described by similar classes of evidence accumulation models^8,57,58^. This observation has led researchers to treat value as a form of evidence that is noisily accumulated in a manner isomorphic to the accumulation of sensory evidence when perceiving a random dot kinematogram. However, given that the dynamics of value accumulation are more difficult to measure and manipulate than the dynamics of perceptual accumulation, questions still remain regarding the basis of value as a form of evidence and the nature of the noise associated with its integration^58^. By manipulating the value associated with sensory evidence accumulated in a multi-attribute decision-making task, the current task could provide leverage in understanding the relationship between these two forms of evidence accumulation. Moreover, the uncertainty our task engenders at the level of both responses and goals (i.e., attentional targets) also makes it well-suited as a potential low-level analog for more complex goal conflicts that occur in daily life, ranging from dietary choice to perseverance on a demanding task in the face of attractive alternatives. While more research is needed to bridge our understanding of how we maintain focus on a paper with our understanding of how we select the words that go on a page, the current findings offer promise that advancing our understanding of one will bring us nearer to closure on the other.

## Methods

### Participants

Thirty-four individuals (71% female, Age: M = 21.1, SD = 2.8) participated in this study. All participants had normal color vision and no history of neurological disorders. Three additional participants were excluded prior to analysis, two due to mechanical errors and one due to an incomplete session. Participants provided informed consent in accordance with the policies of the Princeton University Institutional Review Board.

### Procedure: Choice Task

The main task performed in the scanner required participants to view a random dot kinematogram consisting of red and blue colored dots^3,4^ (Fig. 1). On a given trial, a majority of the dots were either blue or red, and a proportion of the dots (independent of their color) moved in either an upward or downward direction. For consistency with previous studies, we use the term *color coherence* to refer to the relative proportion of red versus blue dots, and *motion coherence* to refer to the proportion of dots moving consistently in one of the two directions. Four coherence levels were used for each attribute, determining varying degrees of discriminability for that attribute on a given trial. For each attribute, these coherence levels were defined as multiples of a single individually-calibrated coherence level that asymptotically produced approximately 80% accuracy on that attribute (see below). For motion, these four levels were 50%, 95%, 140%, and 185% of the calibrated motion coherence level (e.g., if the staircase procedure below settled on a calibrated motion coherence of 10% for a given participant, then the most difficult motion coherence level for this participant would be 5% motion coherence and the easiest level would be 18.5% motion coherence). Initial pilot testing suggested that slightly different scaling values needed to be used for the color attribute in order to more closely match choice preferences across these two attributes, so the equivalent scaling values for the four color levels were 50%, 105%, 160%, and 215% of the calibrated color coherence level. Unless otherwise specified, details of the dot presentation (e.g., color and speed) were identical to Kayser et al (2010), including subjectively isoluminant values of blue and red for the dot colors.

Each color and motion direction was associated with one of two responses (e.g., left button to indicate that the dots are majority blue and/or moving upward; right button to indicate that the dots are majority red and/or moving downward) (Fig. 1A). These response contingencies were counter-balanced across subjects. Participants could only provide one response on each trial (left or right), and this response could be correct for neither, one, or both dimensions. The coherence of each dimension and the congruency across dimensions (i.e., whether or not the same response was correct for both dimensions) was varied independently across trials (Fig. 1B).

Participants were given three seconds (3s) to respond, and the random dot display remained on the screen for that entire duration, including after the response was made. After each trial, participants viewed a fixation cross for 1.2-7.2s (uniformly distributed across trials), which concluded with an additional 0.5s during which the fixation cross changed color to prepare the participant for the onset of the next trial.

Subjects were rewarded based on the number of attributes their (single) response correctly discriminated on a given trial (0, 1, or 2). The rewards for answering each attribute correctly changed over the course of the session, across three epochs of equal length (Fig. 1C): in the first epoch these two dimensions were rewarded equally ($0.15 each); in the second epoch one dimensions was rewarded $0.20 (e.g., motion) and the other $0.10 (e.g., color); and in the final epoch these reward contingencies were reversed (i.e., the attribute that was previously rewarded $0.20 for a correct response was now rewarded $0.10, and vice versa). Each epoch consisted of 192 trials, split across four blocks of 48 trials each.

Before starting the main task, participants performed 16 practice trials outside of the scanner and 16 practice trials inside the scanner (during which no fMRI volumes were collected). These practice trials were followed by feedback on the reward they could have earned for each trial. During the main task (while being scanned) this trial-wise feedback was omitted and participants were only given feedback about average performance at the end of each task block. At the end of the session, 20 trials were selected at random, and participants received the total payment received across those trials.

### Procedure: Psychometric Calibration

Before performing the main task in the scanner, participants performed a task intended to calibrate and match overall performance across the two stimulus attributes. In separate blocks, participants were asked to respond based on one of the two attributes, and the coherence of that target attribute was systematically varied across trials based on a 3-1 psychometric staircase procedure (while the coherence of the alternate attribute was held constant at 0% over that block). Color calibration blocks started at 33% coherence and motion calibration blocks started at 40% coherence. For both block types, coherence was decreased by steps of 1.5% after every 3 consecutive correct trials and increased by the same amount after every error. The participant’s threshold coherence for each attribute was determined based on an average of coherence levels over the last 12 trials of the calibration block.

In order to ensure as stable estimate of the participant’s asymptotic discrimination abilities, the psychometric staircase for each attribute terminated once the following criteria were met: (1) at least 300 trials had passed, (2) the current estimate of threshold coherence (average of coherence levels over the previous 12 trials) was less than 30%, (3) the current estimate of threshold coherence was no greater than 6% (four steps on the psychometric staircase) above the lowest threshold the participant reached over the previous 400 trials (or as many trials had been completed up to that point, whichever were fewer) and (4) there was no significant linear trend in the coherence values over the past 15 trials (i.e., a non-parametric correlation yielded a *p*-value greater than 0.10).

Due to a coding error, the fourth calibration criterion above was not properly implemented for the first seven participants, resulting in coherence thresholds that may have differed slightly from what they would have been assigned with the intended procedure. However, we were unable to find any differences between the behavioral performance of these participants and the remaining participants, in terms of overall accuracy for the high-reward dimension (*z* = 0.1, *p* = 0.89), overall RT (*t* = 1.0, *p* = 0.34), or in the influence of coherence on either choices or RTs (*p*s > 0.48). We therefore include these participants in all of our analyses but note that all of our findings are robust to their exclusion.

### MRI Sequence

Scanning was performed on a Siemens Skyra 3T MR system. We used the following sequence parameters for the main task and localizer: field of view (FOV) = 196mm x 196mm, matrix size = 66 x 66, slice thickness = 3.0mm, slice gap = 0.0mm, repetition time (TR) = 2.4, echo time (TE) = 30ms, flip angle (FA) = 87°, 46 slices, with slice orientation tilted 15° relative to the AC/PC plane. We collected 160 volumes for the decision-making task and 169 volumes for the functional localizers. At the start of the imaging session, we collected a high-resolution structural volume (MPRAGE) with the following sequence parameters: FOV = 200mm x 200mm, matrix size = 256 x 256, slice thickness = 0.9mm, slice gap = 0.45mm, TR = 1.9s, TE = 2.13ms, FA = 9°, 192 slices.

### Behavioral Analysis

All behavioral data were analyzed using mixed-effects regressions in R 3.3.1 (*lmer* and *glmer* functions), modeling all possible subject-wise intercepts and slopes. Response times were log-transformed prior to analysis to reduce skew.

### fMRI Analysis

Imaging data were analyzed in SPM8 (Wellcome Department of Imaging Neuroscience, Institute of Neurology, London, UK). Functional volumes were motion corrected, normalized to a standardized (MNI) template (including resampling to 2mm isotropic voxels), spatially smoothed with a Gaussian kernel (6mm FWHM), and high-pass filtered (128s cut-off period).

Our primary analyses focused on a priori regions of interest (ROIs) within vmPFC, dACC, and areas MT+ and V4 (identified in previous studies and localizers; see below). For these analyses, we generated first-level general linear models (GLMs) that included a separate regressor for each trial, and extracted the associated trial-wise beta estimates for each ROI. These beta estimates were arcsine-transformed (to reduce kurtosis) and then included in mixed-effects regressions across participants, modeling participant-wise random intercepts and slopes.

In order to test for regions sensitive to attribute coherence outside these ROIs, we also performed exploratory whole-brain GLMs. These GLMs modeled event regressors at the onset of each trial (separately for each epoch), with non-orthogonalized parametric regressors for the coherence of each attribute. We then performed second-level analyses consisting of one-sample *t*-tests over contrasts estimated from the first-level GLM. Activations were displayed using a voxelwise *p*-value of 0.005, extent-thresholded to achieve a whole-brain cluster-wise corrected *p* < 0.05.

All first-level GLMs included additional regressors modeling intercepts and linear trends for each task block. Moreover, in order to minimize the influence of outlier time-points (e.g., due to head motion or signal artifact), these GLMs were estimated using a reweighted least squares approach (RobustWLS Toolbox)^59^.

Rather than using the absolute coherence values used for a given participant (e.g., 12%, etc.), fMRI and behavioral regressions coded coherence based on their ordinal levels (1-4). Signed coherence, which reflected the amount of evidence an attribute provided for the response made on that trial, varied from −4 to +4. Positive signed coherence values represented coherence levels that were increasingly consistent with the participant’s response, whereas negative signed coherence values represented coherence levels that were increasingly inconsistent with that response. Unsigned coherence (also referred to as salience), which reflected the overall amount of evidence provided by an attribute irrespective of the response it supports, varied from 1 to 4.

For visualization purposes, plots of beta estimates within a given brain region are based on averaged beta estimates extracted from these whole-brain analyses across subjects. Results of equivalent mixed-effects regressions are reported in the main text.

### Regions of interest

We defined ROIs for dACC and vmPFC based on locations of relevant past findings in research on perceptual or value-based integration of multi-attribute stimuli. Our dACC ROI combined two dACC peaks reported by Kayser et al.^3^, which negatively correlated with the evidence for the attended dimension in a cued-attention version of the current task (MNI coordinates [x, y, z] = 6, 16, 49 and 8, 23, 40; spheres with 6mm radii). Our vmPFC ROI was centered on the peak vmPFC activation from Hare et al.^11^, which positively correlated with the evidence related to two attributes of a value-based stimulus (the taste and health of a food) (3, 36, −12; radius = 5mm). Further analyses within dACC focused on five rostrocaudally arranged 6mm ROIs along the dorsomedial surface, drawn from Taren et al.^33^ (see also Ref. ^38^). This axis ranged from the caudal-most region associated with response conflict (center: −4, 10, 50), to a central region associated with decision conflict (6, 23, 39), to the rostral-most region associated with strategy conflict (-6, 35, 34). Intermediate ROIs fell between the first two ROIs (-4, 16, 45) and the second two ROIs (-4, 30, 37).

### Data availability

All data are available from the authors upon request.

## Acknowledgments

We are grateful to Michael Frank, Harrison Ritz, and Avi Vaidya for feedback on an earlier draft of this manuscript. This work was funded by a postdoctoral fellowship from the CV Starr Foundation (A.S.), and NSF Graduate Research Fellowship (M.A.S.), and by the John Templeton Foundation.

## Supplementary Methods

### Defining motion and color-sensitive regions

We included localizer scans at the end of each session to identify regions that were especially sensitive to dot motion (MT+) and color (V4). Following Kayser and colleagues ^3^, each localizer consisted of ten consecutive 40s blocks, with each block consisting of 10s of increased motion or color information followed by a 30s baseline. For the motion localizer, the 10s period consisted of 100% coherently moving dots, the direction of which changed every second (sampled randomly from the range 0°:36:324°, without replacement), and the 30s baseline consisted of static dots. The color localizer consisted of Mondrian-like images alternating every second in color (10s) versus grayscale (30s baseline).

MT+ and V4 ROIs were generated based on the motion and color localizer tasks, spatially constrained by a priori regions identified by Kayser and colleagues. We performed whole-brain analyses for both localizers, in each case examining regions whose activity was increased during the condition of interest (high-coherence motion or colorful Mondrians) relative to the relevant baseline (static dots or gray patches). The associated first-level analyses and contrasts proceeded as described above. We then used each participant’s localizer to identify the 20 voxels within MT+ (6mm ROI centered at −46, −74, −2) most sensitive to motion and the 20 voxels within V4 (centered at −28, −84, −22) most sensitive to color.

## Supplementary Analyses

### Effect of attribute evidence in MT+ and V4

In spite of their selectivity to motion and color information more generally, previous work explicitly instructing participants to attend either color or motion has found MT+ and V4 show little selectivity for one type of information and instead both negatively track evidence for the attended dimension ^3^. Consistent with these findings, during the biased segments (when reward disproportionately favored one attribute) we found that both regions negatively tracked evidence for the high-reward attribute (e.g., when motion was more rewarding, both MT and V4 negatively tracked signed motion coherence). This was true both when motion (*t_MT_* = −4.0, *t_V4_* = −4.2, *p* < 0.001) and color (*t_MT_* = −2.8, *t_V4_* = −2.1, *p* < 0.05) were the high-reward dimensions; in both cases, these regions did not significantly track the coherence of the low-reward attribute (|*t*s|<1.6, *p*s>0.10). When both attributes were rewarded equally, these regions negatively tracked the coherence of both attributes (motion: *t_MT_* = −2.3, *t_V4_* = −3.7,*p* < 0.05; color: *t_MT_* = −2.2, *t_V4_* = −2.1, *p* < 0.05).

## Supplementary Figures

**Supplementary Figure 1.**
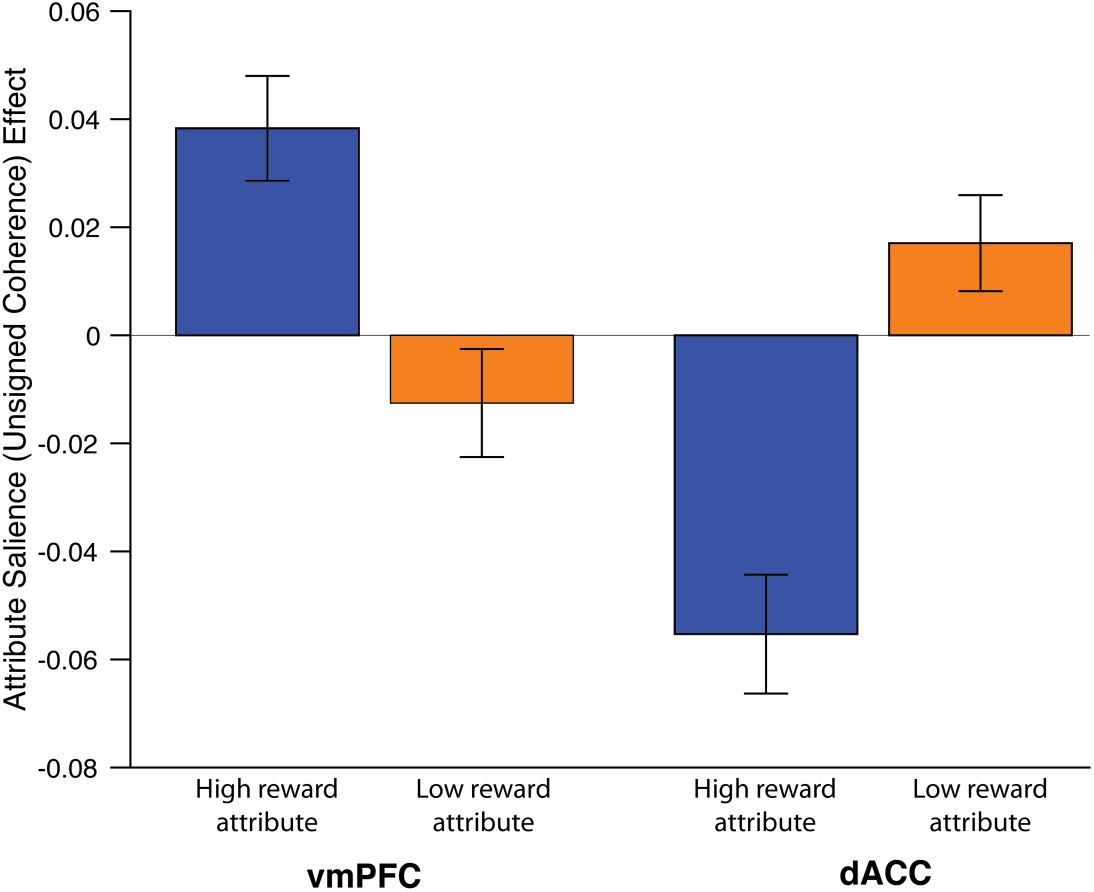
vmPFC and dACC differentially encode the salience of the high and low reward attributes. Whereas vmPFC positively tracked the salience (unsigned coherence) of the high-reward attribute and negatively tracked the salience of the low-reward attribute, dACC exhibited the opposite pattern. Error bars reflect s.e.m.

